# Environmental determinants of round goby invasion refuges at a river scale: implications for conservation of native biodiversity

**DOI:** 10.1101/2022.09.07.506921

**Authors:** Olivier Morissette, Cristina Charette, Matthew J.S. Windle, Abraham Francis, Annick Drouin, Jesica Goldsmit, Alison M. Derry

## Abstract

Introductions of exotic invasive species are a global disturbance for natural habitats. The severity of invasions can greatly vary from local to global scales, as observed in invasion refuges, exhibiting lower-than-expected invasion intensity. In this study, we analyzed the effects of water conductivity and wetland presence on the density of the round goby (*Neogobius melanostomus*) in a large-scale study (> 1300 sites), spanning a 400 km stretch of the St. Lawrence River (Canada). Our results showed that round goby density was null in sites with water conductivity under 100 µS/cm and increased toward a probable biological optimum at 300 µS/cm. The presence of wetlands appeared to also decrease round goby density along the conductivity continuum. Similarly, fish community diversity was maximal outside of the round goby water conductivity optimum. Hence, low water conductivity (<100 µS/cm), in interaction with the presence of wetlands, can provide a refuge for native aquatic species, establishing a simple risk assessment tool for managers. Our results also highlighted the high value of wetland conservation for conservation of native species biodiversity.

## Introduction

Biological invasions are devastating, global anthropogenic disturbances that threaten biodiversity, and disrupt ecosystem functions and services (Bacher et al. 2017; IPBES 2019; Lewis et al. 2016; Pejchar and Mooney 2009; Simberloff et al. 2013). Distressingly, the number of non-native species introductions continues to increase around the world (Seebens et al. 2017; Seebens et al. 2018). The magnitude and frequency of invasions, however, can be highly variable (Dawson et al. 2017). From the global to regional scale, some locations are experiencing a disproportionate invasion pressure and impacts while other neighbouring habitats seem to be far less impacted (Dawson et al. 2017; Dyer et al. 2017; Latzka et al. 2016). This phenomenon is crystallized in the identification of invasion hotspots and refuges, exhibiting contrasting invasion prevalence (number and abundance of invasive species) (Boon et al. 2023; Dawson et al. 2017). Disproportionate levels of invasion across an entire landscape can often be explained by differences in propagule pressure or ecosystem characteristics that mediate invasion success (Catford et al. 2009; Lockwood et al. 2009). However, within invaded ecosystems, other factors that determine whether local sites become invasion hotspots or invasion refuges remain to be better described and understood. Importantly, it has been suggested that spatial environmental heterogeneity (or complexity) and social context (i.e. conservation zones or Indigeneous Peoples presence) within invaded ecosystems seems to play an important role—in conjunction with climate—to promote or suppress invasive species density at local and regional scale and their associated impacts on native species (Melbourne et al. 2007; Schuster et al. 2019; Vander Zanden et al. 2017). Identification and delineation of invasion hotspots and refuges are therefore an effective tool for integrated, multidisciplinary planning of invasive species management and conservation of native biodiversity (Byers et al. 2002).

Invasion refuges could arise within invaded regions or habitats where ecological, social or environmental conditions create the deterrence of invaders, either by decreasing the propagule pressure or the survival or performance of invaders (Anton et al. 2014; Boon et al. 2023; Melbourne et al. 2007; Vander Zanden et al. 2017). The importance of invasion refuges can be crucial as even a small to moderate decrease of invasion pressure or invasive species density can allow coexistence of invasive and native species in neighbouring habitats, hence promoting native species persistence (Bradley et al. 2019; DeRoy et al. 2020; Sofaer et al. 2018; Strayer 2020; Strayer et al. 2018; Yokomizo et al. 2009). The existence of invasive refuges is in accordance with the environmental matching hypothesis, stating that the impacts of an invader decrease with environmental conditions that diverge from its physiological optimum (Iacarella and Ricciardi 2015; Ricciardi et al. 2013). Hence, a growing body of literature has explored environmental factors that promote invasion refuges within invaded freshwater ecosystems (Astorg et al. 2020; Boddy and McIntosh 2021; Boddy et al. 2020; Grabowski et al. 2009; Iacarella and Ricciardi 2015; Kestrup and Ricciardi 2009; Latzka et al. 2016; Priddis et al. 2009). However, many of those studies have not identified the broad role of environmental heterogeneity in the establishment of invasion refuges, mainly because of their limited spatial scales or the context of laboratory experiments (Astorg et al. 2020; Iacarella and Ricciardi 2015; Kestrup and Ricciardi 2009), potentially limiting inferences at the whole ecosystem scale.

The round goby is an invasive littoral benthic fish that was first observed in the St. Lawrence River in 1997 where it became increasingly abundant in many, but not all, habitats (Astorg et al. 2022; Kipp et al. 2012; Morissette et al. 2018). Its geographic range distribution in the St. Lawrence River spans from Lake Ontario to the salt front of the St. Lawrence Estuary, ∼30 km downstream of Québec City. When present, the round goby has strong density-dependant ecological effects, notably by reducing the density of native benthic fishes such as darters (*Etheostoma* spp.) and sculpins (*Cottus* spp.) that use similar habitats (Janssen and Jude 2001; Jude et al. 2018; Morissette et al. 2018; Poos et al. 2009). The round goby can also ravage the diversity and abundance of macroinvertebrate communities (Astorg et al. 2022; Kipp et al. 2012). However, some types of habitats and environmental conditions appear to reduce round goby abundance. In the Great Lakes, low oxygen concentration and low abundance of wetlands or riparian vegetation were linked to invasion success of the round goby in some tributaries (Kornis et al. 2013; Krabbenhoft and Kashian 2022; McAllister et al. 2022). The concentration of dissolved ions (often measured by water conductivity or specific conductance, which is conductivity standardized at 25 °C) seem to also predict the presence and abundance of the exotic round goby in the Great Lakes (Baldwin et al. 2011) and in the St. Lawrence River (Astorg et al. 2020). Another laboratory experiment from a limited portion of the St. Lawrence River showed that the impact of conductivity on round goby performance was influenced by water calcium concentration, where low dissolved calcium ions (Ca^2+^) resulted in low predation rates and, thus, reduced ecological impacts (Iacarella and Ricciardi 2015). Other studies have found evidence to support that wetlands, through their substrate properties and associated submerged aquatic vegetation, offer buffering effects from negative round goby impacts on native biodiversity at invaded sites (Astorg et al. 2020; Cooper et al. 2007). Those results, albeit highly pertinent for risk assessment and management of the round goby, were based on observations from a limited portion of the St. Lawrence River, impeding the potential for a generalization of the results.

The objective of our study was to explore the relative influence of local environmental conditions on the round goby density in a large-scale ecosystem. We used two extensive fish surveys characterizing round goby density and littoral fish community diversity along the entire non-tidal freshwater portion of the St. Lawrence River. Specifically, we aimed to test the hypothesis that variation of water conductivity and presence of wetlands can influence round goby density and native littoral fish diversity, establishing invasion refuges and hotspots. We predicted that non-wetland invaded sites under high water conductivity act as invasion hotspots by increasing round goby density and causing reduced littoral fish diversity. Conversely, we predicted that wetland invaded sites under lower water conductivity would act as invasion refuges from the round goby by reducing round goby density and maintaining or increasing littoral fish diversity. Our study provides information for delimiting round goby hotspots and refuges in the St. Lawrence River, helping to refine risk assessment of Ponto-Caspian invasion in this ecosystem and others under imminent threat of round gobies, as well as promoting the restoration and conservation of invasion refuge in the region.

## Materials and Methods

### Study area

We conducted our study over an area spanning approximately 400 km along the St. Lawrence River, extending from Kingston (Ontario, Canada) at the outflow of Lake Ontario to Batiscan (Québec, Canada) in the St. Lawrence River (Figure 1). The St. Lawrence River is a large, fluvial ecosystem draining the Laurentian Great Lakes toward the Atlantic Ocean. Its narrow fluvial sections successively (upstream to downstream) drain three widening of the St.

**Figure 1.**
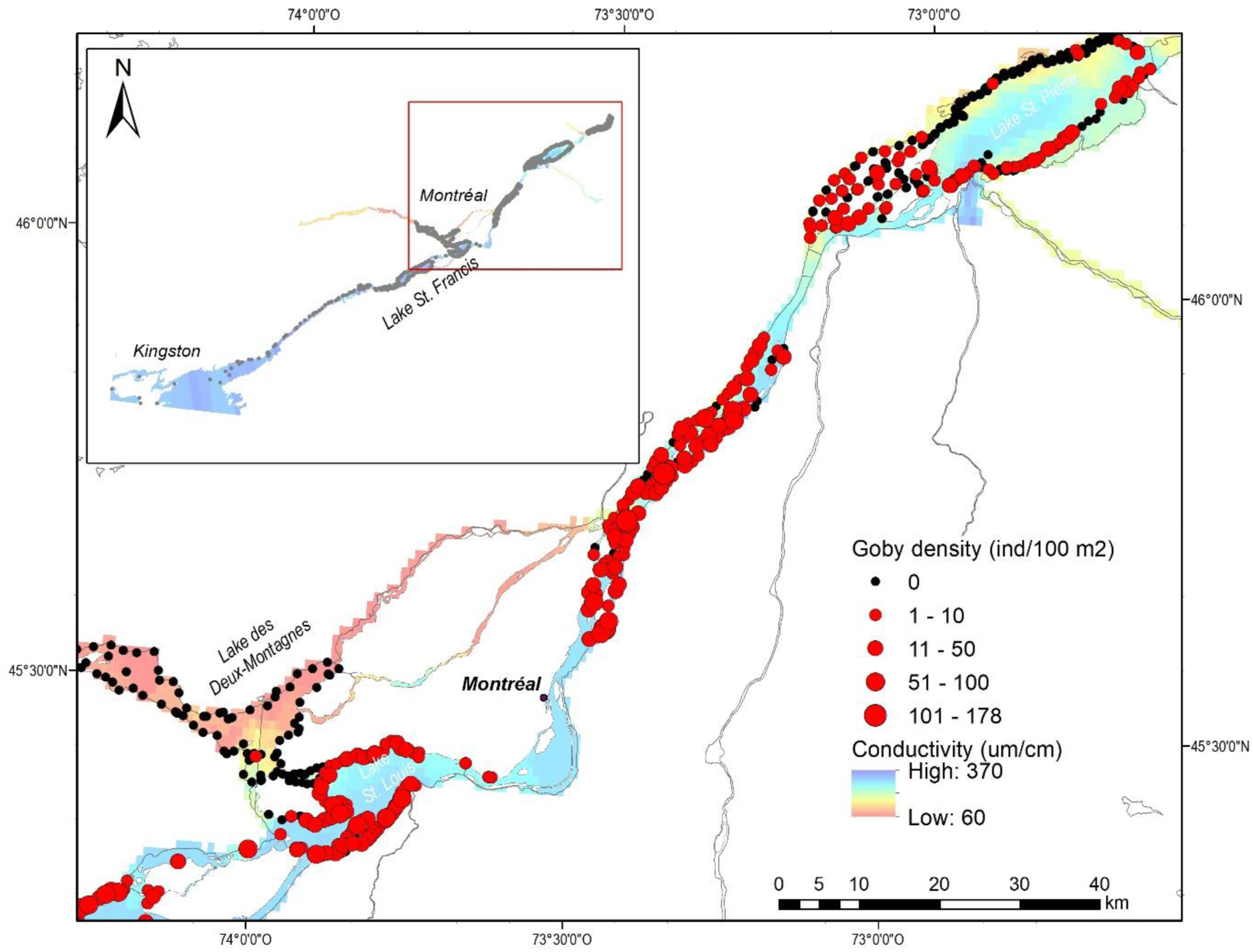
Round goby density (proportional red circles, black circles are null abundance) in relationship to the water conductivity gradient (red to blue colour shade) presented on a subset of the study for visualization (zone of maximum variation). The complete study area is presented in the smaller context map with fish sampling sites locations (gray dots).

Lawrence River, called the fluvial lakes (St. François, St. Louis, and St. Pierre lakes). Where the St. Lawrence reaches the Archipelago of Montréal, the water from the Ottawa River, characterized by its low conductivity (∼60 to 100 µS/cm) and brown coloration, joins the Great Lakes water mass, which has a higher water conductivity (200 to 400 µS/cm) and green coloration. Downstream of Montréal, both water masses flow side by side, with very little lateral mixing (Frenette et al. 2003; Vis et al. 1998), until the St. Lawrence reaches its tidal portion downstream of Lake St. Pierre. Hence, the water masses keep their contrasted chemical properties for hundreds of km, creating a large ecotone where brown waters (Ottawa River) are mostly restricted to the north shore and green waters (Great Lakes) to the south shore. The relative position of the mixing zone and the two water masses stay stable, with seasonal variations depending on the magnitude of spring floods or summer low water. Along the large-scale chemical gradients, various habitats can be observed, from highly vegetated wetlands to sandy and rocky bottom sections, creating an additional environmental variation of interest for fish and invertebrate communities.

### Environmental and fish data

Fish community data were acquired using beach seines as part of both the Fish Identification Nearshore Survey (FINS) conducted by the River Institute and the Mohawk Council of Akwesasne Environment Program, and the *réseau de suivi ichtyologique* (RSI) operated by *ministère des Forêts, de la Faune et des Parcs* fish surveys. The sampling occurred in mid to late summer over 14 years, between July 2007 to September 2021. The FINS fishing was conducted using a 9.1 m long and 1.8 m deep purse seine, operated on three 20 m long hauls for every studied site, corresponding approximately to 182 m^2^ of surveyed habitat per effort. The RSI used a 12.5 m long by 4 m deep beach seine deployed on the site and closed on itself by operators, corresponding to approximately 120 m^2^ of surveyed habitat per site. Both nets were used in coastal, wadable (depth <1.5 m) habitats. Captured fish were identified to species and counted by capture sites. As the spatiotemporal effect of round goby presence on those littoral fish communities have already been assessed (Morissette et al. 2018), we limited the study to the analysis of goby density and fish diversity. Site-specific round goby counts were converted to density (no. gobies/m^2^) to allow comparisons between both FINS and RSI datasets.

Environmental data, conductivity (specific conductance), and wetland presence were either measured and observed on the site during sampling (FINS) or estimated from geographical data available (RSI). During FINS surveys, conductivity (µS/cm) was measured and recorded nearshore (maximum 1-meter depth) at each sampling site using a calibrated water quality multi-parameter sonde (Professional plus model YSI ProDSS). Conductivity estimation was conducted using an independent 1660 sampling points acquired between 2017 and 2021 (MFFP, *unpublished*) spanning all the entire RSI (Québec) sampling area. Conductivity in the RSI sampling zone was interpolated using this dataset and ordinary kriging analysis in the Spatial Analyst toolbox implemented in the software ArcMap 10.7.1 (ESRI). Validation on the interpolation performance was conducted by regression of interpolated and measured conductivity values, showing a reliable agreement (R^2^_adj_ = 0.84, Figure S1). Conductivity values for RSI fish sampling sites were extracted from the interpolated layer using the sampling coordinates and the function *extract* coded within the R package *terra* (Hijmans et al. 2022) implemented in R version 4.1.2 (R Core Team 2021). For both FINS and RSI surveys, presence of wetlands on sites was extracted using two open-access, high-resolution cartography of Ontario (Ministry of Natural Resources and Forestry 2021) and Quebec wetlands (Canards Illimités Canada and ministère de l’Environnement et Lutte contre les changements climatiques 2021). To simplify and correct for wetland definition discrepancy in the datasets, only “open water” class was considered as *non-wetlands* and all remaining classes (e.g., marsh, swamps, wet meadows) were considered as *wetlands*, creating a binary *habitat type* factor.

### Statistical modelling

All statistical analyses were computed in R version 4.1.2 R (R Core Team 2021) and RStudio v2022.02.3+492 (RStudio Team 2020). Generalized additive models (GAMs) were used to assess the effect of conductivity and habitat type on round goby density using the *mgcv* R package v1.8-38 (Wood 2011). GAM is a useful regression-based framework to model non-linear relationships using smooth functions and can incorporate different family error distributions (Wood 2017). GAMs allow for estimation of partial effects of a predictor variable, which represent the weighted sum of multiple basic functions of increasing wiggliness. Three models were built in relation to our hypothesis and predictions. The first two models included round goby density (individual/m^2^) as the response variable. The first model included 4 predictors that were habitat type as a parametric effect, conductivity by habitat factor as a smooth interaction, longitude and latitude as smooth interaction, and year as a random smooth. The second model included all listed predictors, but the conductivity smooth was considered without the interaction with habitat types and its parametric effect (Table S1). For both models, latitude and longitude smooth interaction was included to consider any spatial correlation and to ensure that model residuals are more likely to be independent. Year was identified as a random predictor to generalize the effect over the sampling years and to account for any effect of time since invasion (Astorg et al. 2022). The default thin plate spline was used for the conductivity smooth or for the smooth by factor interaction, the Duchon spline was applied to the latitude and longitude smooth interaction, and a random smooth intercept was used for years. Since the response variable was continuous with left-skewed, non-integer values including zeros, an identity log link function with a Tweedie error distribution was applied to both round goby models.

Model selection was performed on the first two models using the Akaike’s Information Criterion (AIC) score to identify the best-fitted model with the lowest AIC score (Wood et al., 2017) (Table S1). The third GAM included littoral native fish diversity as a response variable. For each site, Shannon diversity index (H) using Hellinger transformed species abundance, without round goby count, was calculated for all sampling sites. The model predictors were also habitat type as a parametric effect, conductivity by habitat factor-smooth interaction, longitude and latitude smooth interaction, and year as a random smooth. The predictors’ splines were identical to preceding models; however, a Gaussian error distribution was applied since response variable (H) was not skewed and normally distributed. Model diagnostics and maximum basis functions (k) were verified using the *gam.check* function and autocorrelation was assessed using the *acf* function of the model residuals from the *stats* R package. No significant autocorrelation was detected in the GAMs and diagnostic plots suggested normality and homoscedasticity of the residuals (Figure S2). (Wood et al. 2017). Model visualization was accomplished using the *mgcViz* package v0.1.9 (Fasiolo et al. 2019) and *ggplot2* package v3.3.5 (Wickham et al. 2016).

## Results

The fish dataset comprised 1310 sampling sites, with an average density of 0.07 goby/m^2^ (±0.17 SD), ranging from 0 to 1.8 goby/m^2^. Conductivity was variable in the study area, ranging from 43 to 576 µS/cm, with an average value of 245 µS/cm (±70 µS/cm SD). The expected spatial structure in water conductivity structure between north shore and south shore of the Upper St. Lawrence River downstream of Montréal was observed (Figure 1), where north shore has a lower conductivity while in the south shore is higher. Measured conductivity in the Upper St. Lawrence was more homogeneous, where variations were mainly influenced by small tributary outflows. Wetlands and non-wetlands were equally represented in the study area, with 662 wetland sites and 648 non-wetland sites in the dataset, with both habitats present in approximately equivalent amounts on both shores (Figure 1). According to our predictions, round goby density showed low abundance from sites of water low conductivity and a sharp increase in median values of water conductivity along this gradient (Figure 2).

**Figure 2.**
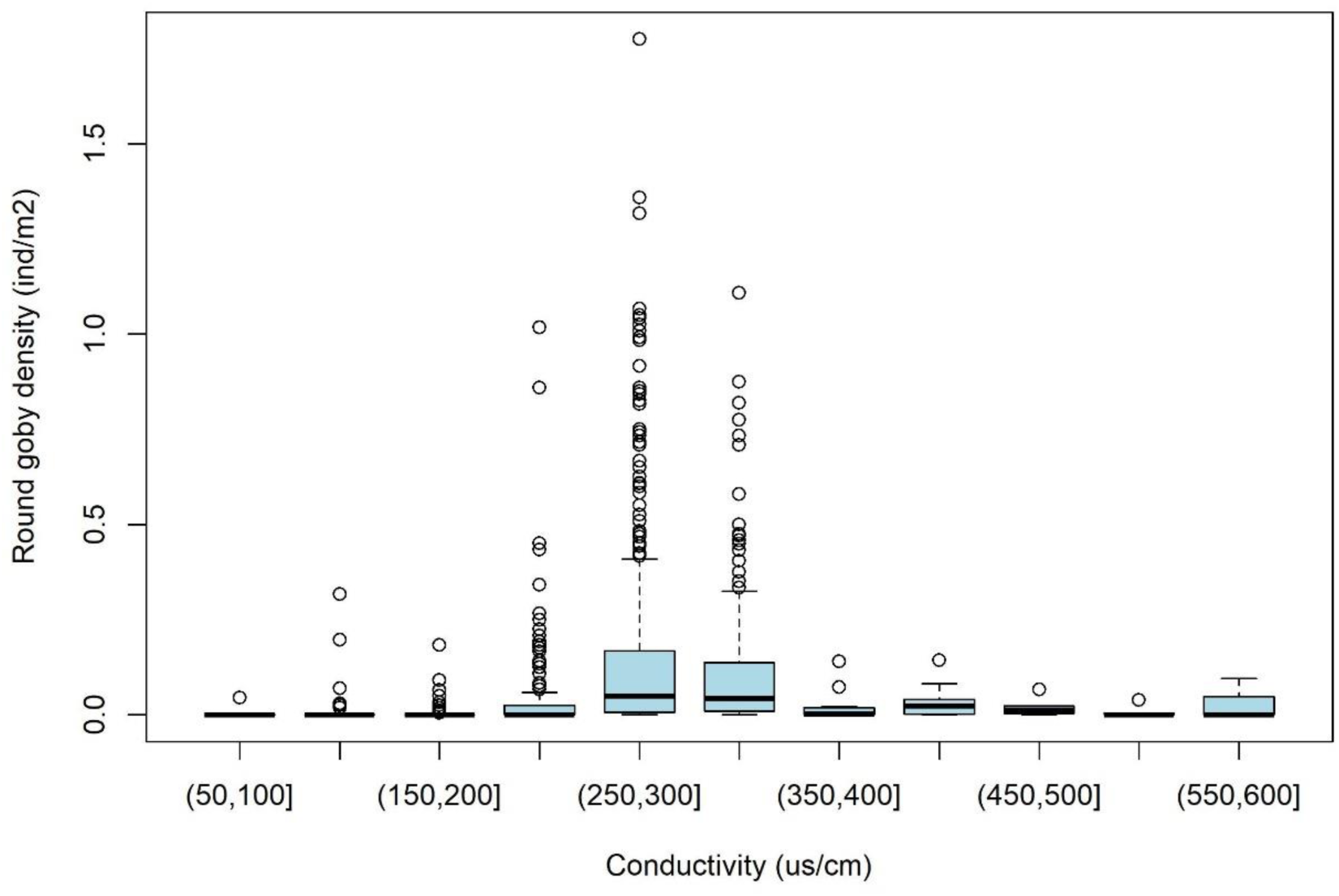
Round goby density (individual/m^2^) by conductivity (µS/cm) for all studied sampling sites. Only 250 to 349 µS/cm groups show a significant difference (ANOVA, F_10,1297_ = 16.73, p < 0.001) in round goby density compared to others.

Our results indicate that both habitat type (wetland/non-wetland) and conductivity were important predictors of round goby density and littoral fish diversity in the St. Lawrence River. The best-fitted GAM model showed the lowest AIC and included habitat type as a fixed effect, conductivity by habitat type factor-smooth interaction along with the latitude and longitude smooth interaction, and year as a random smooth (Table S1 and S2, Figure S2 and S3). A statistical summary of the selected model (Table 1) showed both the fixed parametric effect and partial smooth effects were significant (p-value <0.05) and explained 55.8% of the null deviance with an R^2^-adjusted value of 30.5%. Similarly, the fixed parametric effect of wetlands had a significant (p-value <0.05) on littoral fish diversity, with higher diversity in wetland sites. The GAM explained 21.4% of the null deviance with an R^2^-adjusted value of 18.4%, but the smooth interaction between conductivity and wetlands did not produced a significant effect, only showing a trend of non-wetlands interaction (p = 0.07).

**Table 1.** Statistical summary of the generalized additive model (GAM) selected including habitat type as a fixed effect, conductivity by habitat type factor-smooth interaction, longitude and latitude smooth interaction, and year as a random smooth. Edf indicates the estimated degree of freedom and Ref.df represents the residual degree of freedom. Significant parametric (A) and partial effects (B) are indicated by p-value <0.05.

| A. parametric coefficients | Estimate | Standard Error | t-value | p-value |
| --- | --- | --- | --- | --- |
| (Intercept) | -3.8625 | 0.1487 | -25.97 | <0.0001 |
| Wetlands | -0.4896 | 0.1479 | -3.31 | <0.005 |
| B. smooth terms | edf | Ref.df | F-value | p-value |
| s (conductivity): non-wetlands | 7.198 | 8.941 | 4.127 | <0.0001 |
| s (conductivity): wetlands | 4.15 | 5.246 | 6.764 | <0.0001 |
| s (longitude, latitude) | 55.791 | 79 | 17.365 | <0.0001 |
| s (year) | 9.85 | 14 | 3.003 | <0.0001 |
| R2-adjusted | 0.305 |  |  |  |
| Deviance explained | 55.8 % |  |  |  |

Habitat type showed a direct effect on round goby density and littoral fish community diversity. Wetlands generally hosted lower round goby density and preserved a higher fish diversity compared to non-wetlands (Figure 3). Water conductivity showed a non-linear, partial effect on the round goby density and diversity. However, the magnitude of the effect was different between habitat types and was dependent on the range of the observed values. Overall, from 40 µS/cm to approximately 220 µS/cm, conductivity showed a negative effect on round goby density (Figure 3). This negative effect was found to be amplified in wetlands compared to non-wetland habitats (Figure 3). Above this threshold, water conductivity displayed a positive effect on round goby density in both habitats. At higher levels of conductivity (> 380 µS/cm), a negative effect on round goby density was only observed in wetlands. However, the confidence interval is large and includes the null effect. Overall, the round goby exhibited a maximum density around 295 µS/cm, a potential biological optimum. Accordingly, fish community diversity increased from 40 µS/cm to 190 µS/cm, but declined as round goby density increased (220 to 300 µS/cm), to reach its minimum around the round goby optimum (295 µS/cm). The confidence intervals were also large and included the null effect.

**Figure 3.**
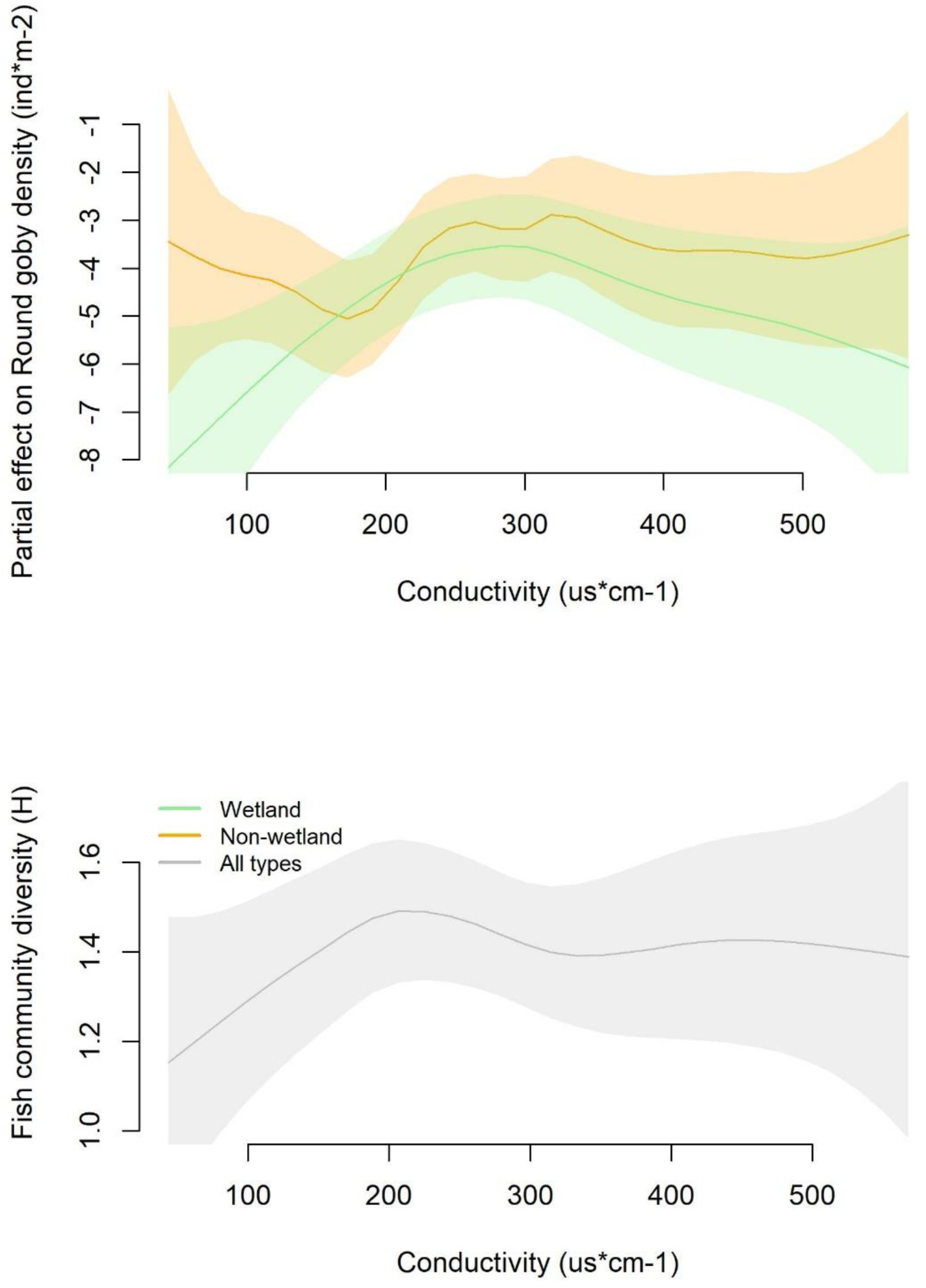
Generalized additive model (GAM) partial smooth effect of conductivity (µS/cm) on round goby density (top, individual/m^2^) and littoral fish community diversity (bottom, Shannon H) by habitat types: non-wetlands (orange), wetlands (green) or all types (gray, showing non-significant interaction). Shaded area indicates the 95% confidence interval.

The best-fitted GAM model demonstrated a significant partial effect of latitude and longitude on round goby density (Figure 4), showing a spatially coherent weighted sum of models’ predictors on round goby density. A considerable portion of the south shore of the St. Lawrence River displayed an important positive effect on round goby density while some locations, notably the Ottawa River and Deux-Montagnes Lake (a widening of the Ottawa River), had the most negative partial effect on round goby density. We consider that all the acquired information could guide round goby risk assessment in a simple two-metrics fashion (Table 2), considering both conductivity and wetland presence. Herein, we propose that non-wetland habitats with water conductivity under 100 µS/cm have a very low risk of goby impacts, evaluated as nearly null in wetlands under this conductivity. This risk increases with increasing water conductivity until higher than 300 µS/cm, reaching the ecological optimum where impacts are predicted to be maximal. Freshwater habitats over this threshold are known to host round goby populations with substantial impacts (Baldwin et al. 2011; Kipp et al. 2012; Kornis et al. 2013). However, under these conductivity conditions, wetlands still have buffering effects, notably by preserving littoral fish community diversity.

**Figure 4.**
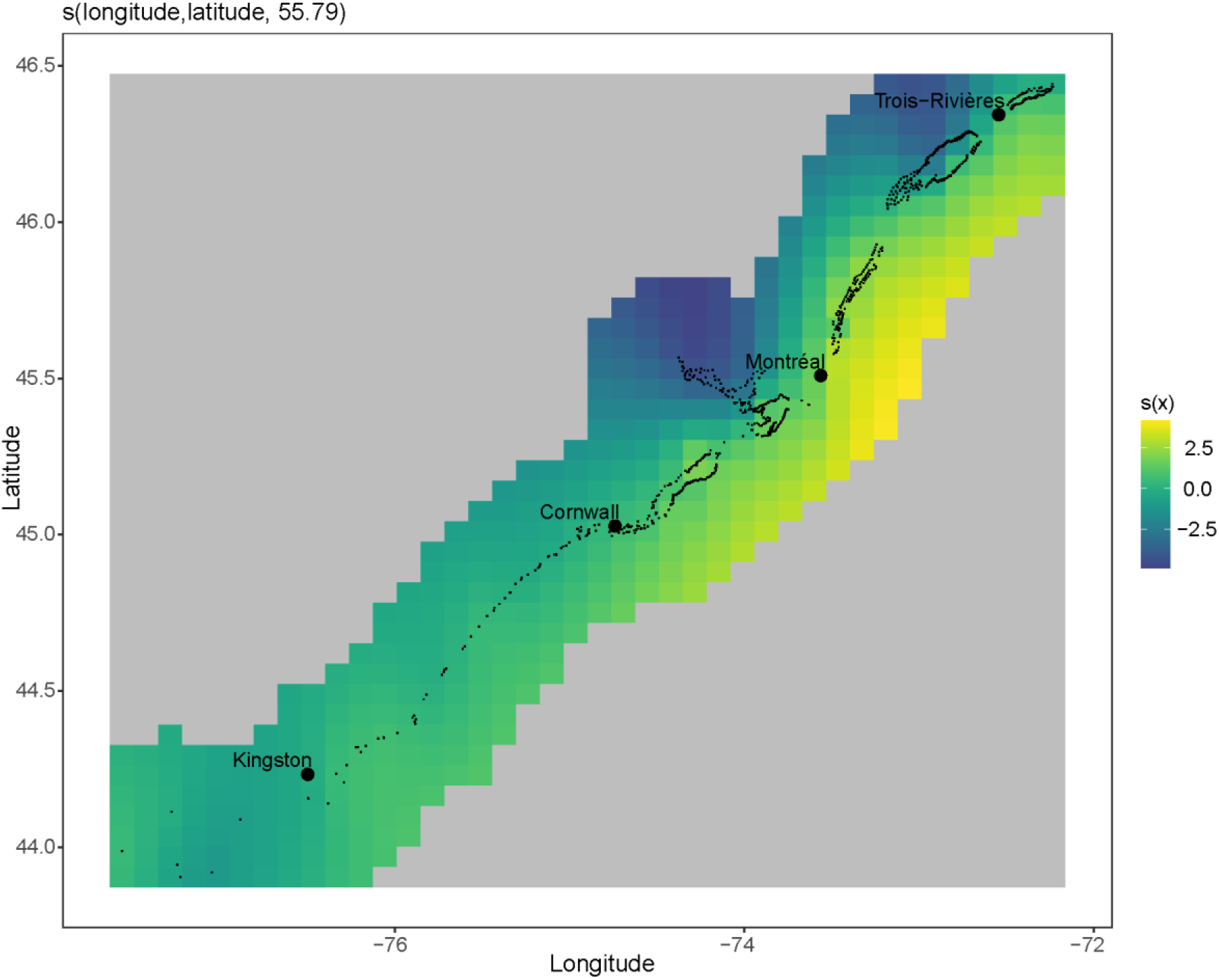
Generalized additive model (GAM) of partial spatial smooth effect on round goby density (individual/m^2^) along the geographic gradient of the St. Lawrence River study area. The range of colours represents the effect: yellow shows the most positive effect (higher density) and dark blue the most negative effect (lower density). Partial residuals are represented by dots.

**Table 2.** Binary risk of round goby density (and density-dependent impacts) evaluation grid based on specific conductance (conductivity) and local presence of wetlands on a local to large-scale study site.

| Local habitat | Water specific conductance (conductivity) |  |  |  |
| --- | --- | --- | --- | --- |
| | <100 $\mu\text{S}/\text{cm}$ | 100 to 200 $\mu\text{S}/\text{cm}$ | 200 to 300 $\mu\text{S}/\text{cm}$ | > 300 $\mu\text{S}/\text{cm}$ |
| Non-wetlands | very low | low | medium | high |
| Wetlands | none | very low | medium* | high* |

\* General agreement to large-scale risk with possibility of local or seasonal buffering effects of wetlands

## Discussion

Our study aimed to examine the effects of two environmental conditions—water conductivity and presence of wetlands—on the invasive round goby density and littoral fish community diversity at a large-scale ecosystem, the St. Lawrence River. This study represented a test of the hypothesis stating that those condition could establish large-scale and local invasion refuges and hotspots. Using two unpublished and extensive fish survey datasets, we applied GAMs to determine thresholds for ecological optimum along the water conductivity gradient and showed large-scale effects of water conductivity on round goby density, as previously recognized in its invaded range (Astorg et al. 2020; Baldwin et al. 2011). Our results revealed that round goby density was positively influenced by increasing conductivity until reaching a probable biological optimum of ∼300 µS/cm (invasion hotspots). Sites of water conductivity under 200 µS/cm had low density to complete absence of round gobies (<100 µS/cm), even in regions hosting established populations for more than 10 years (Morissette et al. 2018). These sites therefore represent possible invasion refuges, a hypothesis supported by the higher diversity observed in those sites. Additionally, even if the presence of wetlands did not completely disrupt the pattern of water conductivity effects on round goby density in invaded habitats, wetlands appeared to decrease round goby density and preserve littoral fish community diversity along the conductivity continuum, contributing to the invasion refuge effect. Our study addressed the influence of those two environmental factors along an over 400 km stretch of the Upper St. Lawrence River, compiling more than 1300 capture sites on a 14-year-old-time series. Hence, our results provide an empirical criterion delineating round goby invasion hotspots and refuges. The sheer size of this study underlines the possibility of generalizing the phenomenon and emphasizes the importance of habitat diversity (i.e., wetland conservation and restoration) to buffer the effects of biological invasion.

The detection of a predictable response of round goby density in the Upper St. Lawrence River suggests that the risk of potential impact, that we consider density-dependent, could be identified by two simple metrics, water conductivity and wetlands presence, participating in identification of sites likely to be more or less impacted by round goby presence. Hence, wetlands and low conductivity could represents significant contribution to establishment of ecological refuges from the round goby, even if these habitats characteristics can be limited in spatiotemporal scale, depending on submerged vegetation abundance, density and phenology (Janetski and Ruetz 2015; Seilheimer and Chow-Fraser 2007). The simple risk grid that our study provides can offer a fast and an easy-to-implement tool for managers and could be disseminated on a large landscape with relatively accessible data. Management of aquatic invasive species is often confronted with the challenge of rapidly identifying the present or upcoming threat of a given species which can cause a vast array of impacts (Lodge et al. 2006). Identification of those impacts can largely vary in the invaded range and depart from what has been identified in laboratory experiments and small-scale field studies. A broader understanding of context-dependent potential density and their related impacts is needed to implement an efficient management in heterogeneous landscapes and populate a more refined impacts evaluation framework (Vander Zanden et al. 2017). Our finding that a local-scale phenomenon, observed in 120 m^2^ habitat patches, had a uniform response across a large environment scale underscores the value of observational studies in invasion biology and its unique potential to offer generalizable risk assessment tools.

The negative effect of low water conductivity on round goby density is concordant with field observations made by Astorg et al. (2020) and Baldwin et al. (2011), and our findings add to a known effect of low conductivity on Ponto-Caspian invaders (Kestrup and Ricciardi 2009; Zhulidov et al. 2004). Reduced Ponto-Caspian invader density in waters with low concentrations of dissolved ions are suggested to be linked with the evolutionary history of these euryhaline species, which experienced repeated marine invasions—characterized by extreme variations of salinity—favouring physiological processes requiring availability of high-dissolved ion concentrations in ambient waters. Differences in round goby performance along freshwater conductivity gradients may be linked to the concentration of calcium ions (Ca^2+^), which contribute to physiological processes and represent a likely common element between freshwater and saltwater (Baldwin et al. 2011; Hunn 1985; Iacarella and Ricciardi 2015; Mayer et al. 1994). Calcium represents a critical component of homeostasis in fish, by intervening on rates and efficiency of gill-water exchanges. Accordingly, field observations suggest that the fish calcium uptake from the environment seems to represent an evolutionarily conserved process that does not proportionally respond to large differences in environmental calcium, even after long-term (multiple generations) residence along this environmental gradient (Sanderson et al. 2023). Hence, the poor performance of round gobies—along with other Ponto-Caspian invaders—can be a manifestation of the limited plasticity of this trait under the unusual environmental conditions imposed by low conductivity tributaries or water masses. This is an eloquent example of environmental matching hypotheses and how the intolerance for low conductivity waters translates into natural refuge to invasion (Ricciardi et al. 2013).

The mitigation of round goby presence and density in the St. Lawrence River by environmental heterogeneity in water conductivity and local presence of wetlands could represent an important factor for the resilience of native fishes and macroinvertebrates. The resistance of wetlands to round goby invasion was previously identified within the St. Lawrence and Great Lakes Basin (Astorg et al. 2020; Cooper et al. 2007), but the mechanisms remain to be identified. Cooper et al. (2007) suggested that such a phenomenon could be linked to time since invasion or lack of access to rocky and hard substrates, a consideration also considered by Kornis et al. (2013). Our study, however, shows that round goby invasions are severe at local sites with favourable environmental conditions and remain absent in unfavourable sites, even if the establishment occurred 2 decades ago (Astorg et al. 2022; Morissette et al. 2018). This refutes the first hypothesis and supports Krabbenhoft & Kashian’s (2022) conclusions that sites showing riparian degradation may be more subject to invasion and their impacts. Although our models did not consider local substrate (non-available), rocky and boulder banks are marginal in the studied region (Sergy 2008), suggesting that substrate may be of minor influence on observed density in and out of the wetlands. Hence, we suggest that factors promoting invasion resilience of wetlands may be linked to the impediment of movements caused by submerged vegetation, predation pressure or different feeding opportunities for the round goby (e.g. dreissenids mussels presence or macroinvertebrates); these are research avenues that remain to be tested.

Aside from its role buffering invasion impacts identified in our study, wetlands in freshwater habitats are contributing to a vast array of ecological functions and services and are of great importance for indigenous peoples living at their proximity (i.e., medicine cabinet and sources of food). The sum of those ecosystem services and socioeconomics benefits underlines the critical importance of wetlands in aquatic habitats. Sadly, wetlands have experienced drastic declines, disappearing three times faster than forested areas (Convention on wetlands 2021). Such declines, fuelled by anthropogenic modifications of freshwater habitats, pollution and climate change (Johnson et al. 2010; Patenaude et al. 2015), can still be contained and even reverted with the application of emergency plans of conservation (Tickner et al. 2020). Based on our results, we argue that wetlands can also be viewed as a pragmatic tool in the integrated management of water quality and prevention of biological invasions (Astorg et al. 2020; Patenaude et al. 2015; Trepel 2010). Promotion of a diverse, yet complex mosaic of heterogeneous habitats could potentially decrease invasion impacts and offer refuge for native species, including the imperiled species susceptible to round goby impacts, notably the eastern sand darter (*Ammocrypta pelludica*), the channel darter (*Percina copelandi*) and the pugnose shiner (*Notropis anogenus*), thus favouring their coexistence and persistence (Melbourne et al. 2007; Priddis et al. 2009). Additionnally, conservation of wetlands along the St. Lawrence River have been important for many indigenous communities, which may have established critical invasion refuges in their surrounding (Schuster et al. 2019), however, our actual dataset cannot serve to test this hypothesis.

We recognize that factors for invasion success and ecological impacts of biological invasions are complex and subject to multiple—and potentially synergistic—biotic and abiotic influences (Catford et al. 2009; Ricciardi et al. 2021; Spear et al. 2021). Notably, over the large study region, the influence of climatic variations (i.e., water temperature, amount of precipitation, winter severity) could also contribute to the observed spatial effect. However, contrasted round goby density observed in close sampling sites (i.e., in the vicinity of Montréal and within Lake St. Pierre) suggest that climatic variations were probably of low importance in the observed patterns that we report here. Round goby invasion success could be influenced by other factors that were not addressed in our study, such as the importance of propagule pressure, feeding opportunities (i.e., abundance of dreissenid mussels or invertebrates), other environmental variables (i.e., sediment type and water flow), or ecological relationships (i.e., predation). We also recognize that maximum density observed in the St. Lawrence River (1.8 goby/m^2^) remains lower than the maximum observed in the Great Lakes, reaching at least 3.5 goby/m^2^ (Taraborelli et al. 2009), which raises questions of other underlying factors contributing to round goby density. Overall, however, the striking effect of water conductivity on round goby density, and therefore ecological impact on native fish and macroinvertebrate communities, offers an interesting avenue to study mechanisms under invasion success and magnitude.

In conclusion, our study offered a unique opportunity to combine two ongoing and long-term systematic fish surveys along a highly structured and complex freshwater ecosystem. We observed a consistent effect of water chemistry and wetland presence in decreasing the density of a notable Ponto-Caspian invader. Our results showed that low water conductivity (<200 µS/cm), in interaction with the presence of wetlands, is linked to a lower round goby density, buffering the potential ecological impacts of this invader and providing a refuge for native aquatic species such as macroinvertebrates and fish. The combination of easily measurable environmental metrics can provide a simple, yet robust risk assessment tool for managers. Our results also highlight the high value of wetland conservation and restoration— led by a diverse community of peoples practicing various uses—as a promoter of biodiversity and an integrated management tool for conservation of native species, especially in invaded ecosystems subjected to anthropogenic disturbances.

## Acknowledgments

The authors are grateful for the field crew and partners who contributed to the FINS and RSI, notably Chantal Côté, Yves Paradis, Kate Schwartz, and staff at the Mohawk Council of Akwesasne’s Environment Program. All fish collections were approved under the Institutional Animal Protection Committee either at MELCCFP, UQAM, and the River Institute and authorized by licences for scientific fish collections from the Ontario Ministry of Natural Resources and Forestry and the Ministère de l’Environnement, de la Lutte contre les changements climatiques, de la Faune et des Parcs du Québec.

## Data availability statement

All raw data files and R scripts used in this work are freely available in the following Borealis repository [<u>Borealis link will be provided upon acceptance</u>].

## Authors’ contributions

All authors contributed to the ideation of the study, the manuscript writing and the revision. MW and CC contributed to the FINS survey and OM and AD contributed to the RSI survey. MW, CC, and OM did the data curation and quality assessment, while CC and OM performed the statistical analyses and produced figures and tables.

